# Structural basis of transcriptional activation by the OmpR/PhoB-family response regulator PmrA

**DOI:** 10.1101/2022.07.20.500760

**Authors:** Yuan-Chou Lou, Hsuan-Yu Huang, Hsin-Hong Yeh, Wei-Hung Chiang, Chinpan Chen, Kuen-Phon Wu

## Abstract

PmrA, an OmpR/PhoB-family response regulator, activates gene transcription responsible for polymyxin resistance in bacteria by recognizing promoters in which the canonical *-35 element* is replaced by the *pmra-box*, representing the PmrA recognition sequence. Here, we report a cryo-electron microscopy-derived structure of a bacterial PmrA-dependent transcription activation complex (TAC) containing a PmrA dimer, an RNA polymerase σ70-holoenzyme (RNAPH), and the *pbgP* promoter DNA. Our structure reveals that the RNAPH mainly contacts the PmrA C-terminal DNA binding domain (DBD) via electrostatic interactions and reorients the DBD three base pairs upstream of the *pmra-box*, resulting in a dynamic TAC conformation. *In vivo* assays show that substitution of PmrA DNA-recognition residues eliminated its transcriptional activity, but variants with altered RNAPH-interacting residues exhibited elevated transcriptional activity. Our study indicates that both PmrA recognition-induced DNA distortion and PmrA promoter escape play important roles in its transcriptional activation.

## INTRODUCTION

PmrA is the primary response regulator of genes responsible for polymyxin resistance in various bacterial species, such as *Salmonella enterica, Escherichia coli*, and *Klebsiella pneumoniae*^1-3^. The PmrA binding promoter, termed the *pmra-box* and consisting of two hexanucleotide XTTAAY repeats separated by five nucleotides, overlaps with the *-35 element* in the *pbgP, ugd*, and *pmrCAB* operons^2,4,5^. PmrA binding and activation of these genes encoding enzymes that modify lipopolysaccharide in the outer membrane promotes resistance to polymyxins, antimicrobial peptides, and antibiotics^2,5,6^. Polymyxins, especially polymyxin B and colistin (polymyxin E), are considered last-resort antibiotics against multidrug-resistant Gram-negative bacteria^7,8^. However, polymyxin-resistant infections are becoming widespread, representing a significant threat to public health^9,10^. Thus, understanding the structural basis of the PmrA-modulated transcriptional activation that promotes bacterial polymyxin resistance is urgently needed to aid the development of new drugs against bacterial infections.

PmrA belongs to the OmpR/PhoB family of response regulators, comprising an N-terminal receiver domain (REC) and a C-terminal winged-helix DNA binding domain (DBD) that are connected by a flexible linker^11^. For PmrA and several other family members, phosphorylation-induced dimerization of the REC allosterically enhances binding of the DBD to promoter DNA for transcriptional activation^12-14^. Despite structural similarities, OmpR/PhoB family members display different modes of interaction with RNA polymerase (RNAP) to regulate transcription. Mutational studies have suggested that OmpR operates by its DBD interacting with the α subunit of RNAP^15,16^, whereas the PhoB DBD interacts with the σ4 domain of RNAP σ70-holoenzyme (RNAPH)^17-19^. A low-resolution crystal structure of a PhoB subcomplex has been reported previously that revealed interactions between a chimeric σ4 domain of the σ70 RNAP cofactor, two PhoB DBDs, and the promoter DNA^20^. However, this subcomplex could not illustrate the full suite of interactions between PhoB and RNAPH. Moreover, a previous attempt to resolve a transcription activation complex (TAC) involving the OmpR/PhoB-family response regulator KdpE by cryogenic electron microscopy (cryo-EM) proved unsuccessful^21^, with the density map for KdpE missing even though the KdpE-RNAPH-DNA ternary complex had been cross-linked with glutaraldehyde. Thus, further structural analyses are required to reveal all interactions between an OmpR/PhoB family member and the RNAPH.

Previously, we found that beryllofluoride (BeF_3_^−^, a phosphate analog)-activated PmrA forms a dimer displaying enhanced binding affinity for the *pbgP* promoter^22^. We also determined a 3.2-Å– resolution crystal structure of BeF_3_^−^-activated PmrA-WI (a W181G/I220D double-substitution variant that exhibits high solubility and thermal stability) in complex with the *pbgP* promoter DNA^14^. In that structure, the two REC domains form a head-to-head dimer, but the two DBDs bind to the DNA in a head-to-tail orientation, resulting in an asymmetric REC-DBD interface in the upstream PmrA protomer, which was demonstrated to be transiently populated and was not critical for transcriptional activation. Our complex structure also revealed details of PmrA-DNA interactions, with the two DBDs specifically recognizing repeated “TAA” sequences in the *pbgP* promoter and inducing a bent conformation in the promoter DNA. Furthermore, a β-galactosidase reporter assay demonstrated that the W181G/I220D double-substitution did not affect PmrA’s transcriptional activity, but the DNA-recognition residues are crucial for PmrA-activated gene expression. In this study, we determine the structure of an intact PmrA TAC ternary complex containing *K. pneumoniae* PmrA-WI, *E. coli* RNAPH, and the *K. pneumoniae pbgP* promoter. The resulting cryo-EM map shows that contacts between the PmrA dimer and its surrounding area are dynamic, and the respective densities could be further enhanced by 3D conformational variability analysis. The final TAC structure reveals interactions between full-length PmrA dimer, RNAPH and DNA. Combined with *in vivo* β-galactosidase reporter assay data, we propose a mechanism for PmrA-mediated transcriptional activation.

## RESULTS AND DISCUSSION

### Overall structure of the PmrA-dependent TAC

In this study, we constructed a synthetic DNA scaffold of the PmrA-dependent promoter (from - 62 to +14 base pairs, 76 bp in total), which contains an upstream AT-rich sequence and the *pmra-box* from the *K. pneumoniae pbgP* promoter (-62 to -13 bp)^2^, a consensus *-10 element*, a discriminator element, a transcription bubble and a 14-bp downstream extension (Figure S1A). The synthetic promoter was mixed with BeF_3_^−^-activated PmrA-WI and the *E. coli* RNAPH for cryo-EM grid preparation (Figure S1B). Sequences of *K. pneumoniae* and *E. coli* RNAPH are almost identical (99%, 98% and 98% for the α, β and β’ subunits, respectively, and 96% for the σ70 cofactor), implyinga strong likelihood of their reasonable hybridization to the PmrA-dependent TAC. We constructed a cryo-EM map of the PmrA TAC at a resolution of 3.03 Å using a single-particle reconstruction approach and advanced 3D conformational variability analysis^23^ (Figure S2). Our map shows a complex of RNAPH, upstream DNA, a PmrA dimer (denoted PmrA-1 and PmrA-2 for the upstream and downstream PmrA protomers, respectively), and the non-conserved region of σ70, revealing the structure of RNAP core subunits and the region of DNA spanning base pairs -42 to 14, including the transcription bubble (Figure 1B). Local resolution estimation of this cryo-EM map indicated that the RNAPH features a static state relative to the dynamic coupling of PmrA and the upstream DNA region (-65 to -43 bp) (Figure S2). Our map also reveals a PmrA-bound transcription activation machinery. We then completely reconstituted the structure of the PmrA TAC based on both maps (Figure 1). The detailed data collection parameters and refinement statistics are summarized in Table 1.

**Table 1.** Cryo-EM data collection, refinement and validation statistics.

|  |  |
| --- | --- |
| Accession codes (PDB/EMDB) | 7W8C / 32354 |
| **Data collection and processing** |  |
| Microscope/detector | Krios/K2 |
| Magnification | 165,000 |
| Voltage | 300 kV |
| Electron exposure (e-/Å<sup>2</sup>) | 57.1 |
| Defocus range (μm) | -1 to -2.2 |
| Pixel size (Å) | 0.822 |
| Symmetry imposed | C1 |
| Initial particle images (no.) | 1,113,892 |
| Final particle images (no.) | 55,568 |
| Reconstruction method | Single particle reconstruction |
| Map resolution (Å) (FSC<sub>0.143</sub>) | 3.03 |
| Map resolution range (Å) | 2.0-8.8 |
| Map sharpening B factor (Å<sup>2</sup>) | 49.4 |
| **Refinement** |  |
| Initial model | 4S04, 6BC6, 6B6H and 6CA0 |
| **Model composition** |  |
| Non-hydrogen atoms | 36353 |
| Protein residues | 4231 |
| Nucleotides | 152 |
| **B factor (Å<sup>2</sup>)** |  |
| Protein | 71.61 |
| Nucleotides | 191.55 |
| **R.m.s. deviations** |  |
| Bond lengths (Å) | 0.010 |
| Bond angles (°) | 1.185 |
| **Validation** |  |
| MolProbity score | 2.2 |
| Clashscore | 15.89 |
| Poor rotamers (%) | 1.00 |
| **Ramachandran plot** |  |
| Favored (%) | 91.63 |
| Allowed (%) | 8.26 |
| Disallowed (%) | 0.12 |

**Figure 1.**
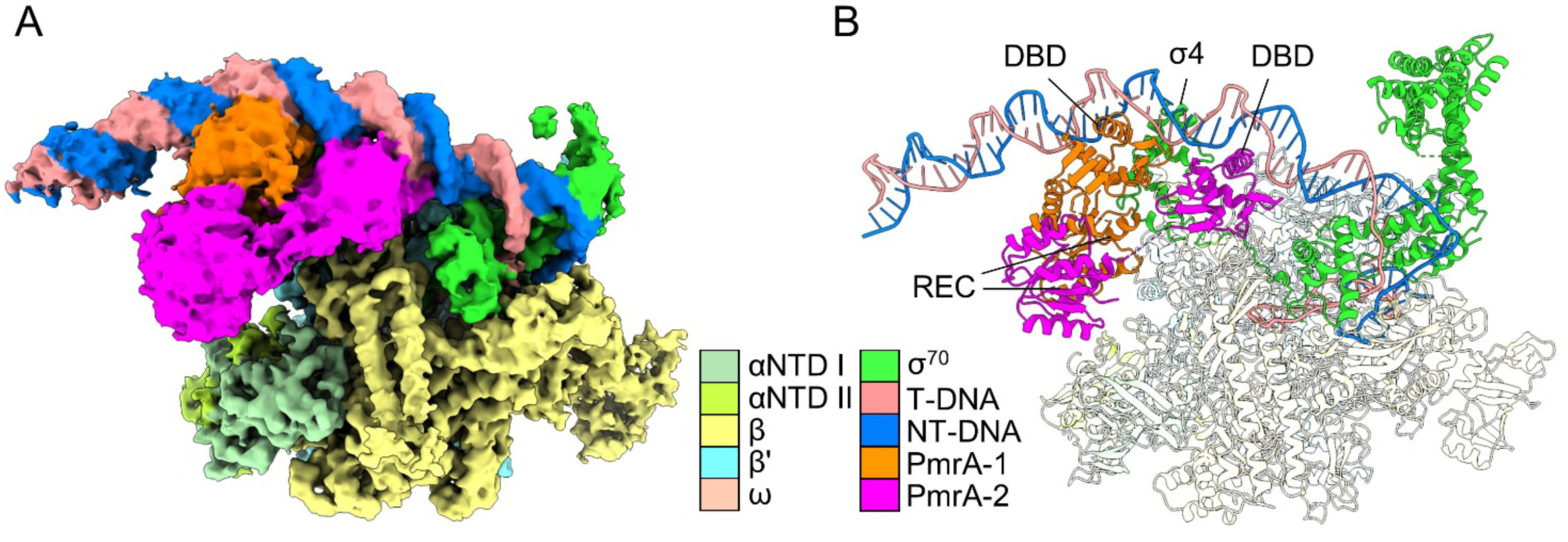
Cryo-EM structure of the PmrA-dependent transcription activation complex (TAC). (A) Cryo-EM map of the PmrA TAC. The PmrA dimer and upstream DNA have been enhanced using the 3D Variability Analysis algorithm. Protein and DNA components are color-coded (see legend). PmrA-1 and PmrA-2 denote the upstream and downstream PmrA protomers, respectively. The double-stranded DNA and PmrA dimer are shown in blue/salmon and orange/magenta, respectively. (B) PmrA TAC structural model displayed in contrast-enhanced cartoon mode, with double-stranded DNA, PmrA dimer and the σ70 cofactor colored as in (A). RNAP core proteins are colored white.

### Electrostatic interactions between PmrA and the RNAPH

In our PmrA TAC, the *E. coli*-derived RNAPH adopts an overall conformation similar to that displayed by the *E. coli* RNAP σ70 transcription initiation complex (TIC)^21^, and the conformation adopted by the PmrA dimer is similar to when it is in complex with DNA alone (Figure S3), demonstrating that the RNAPH and PmrA conformations largely do not interfere with each other. First, we investigated the interactions between PmrA and RNAPH and found that the two PmrA protomers are in close contact with different regions of the RNAPH (Figure 2). PmrA-1 interacts with the σ4 domain and β-flap via its DBD and REC, with an interface of 650 Å^2^. The DBD of PmrA-2 contacts the β-flap and β’ subunit, with an interface of 515.7 Å^2^. Although the resolution of our cryo-EM data does not allow precise characterization of the interacting atoms, we inferred the electrostatic interactions based on the amino acids with opposing surface charges at appropriate distances. An acidic patch (E172, D182, and E184) on the PmrA-1 DBD faces a basic patch (K593, R596, K597 and R599) of σ4 (Figure 2A). Residue D182 of the PmrA-2 DBD also forms electrostatic interactions with K900 at the β-flap-tip helix and with R77 on the β’ subunit (Figure 2B). Furthermore, two additional electrostatic interactions—between R68 on the PmrA-1 REC and E859 on the β-flap and between R160 on the PmrA-2 DBD and D912 on the β-flap—further secure PmrA to the β subunit of the RNAPH.

**Figure 2.**
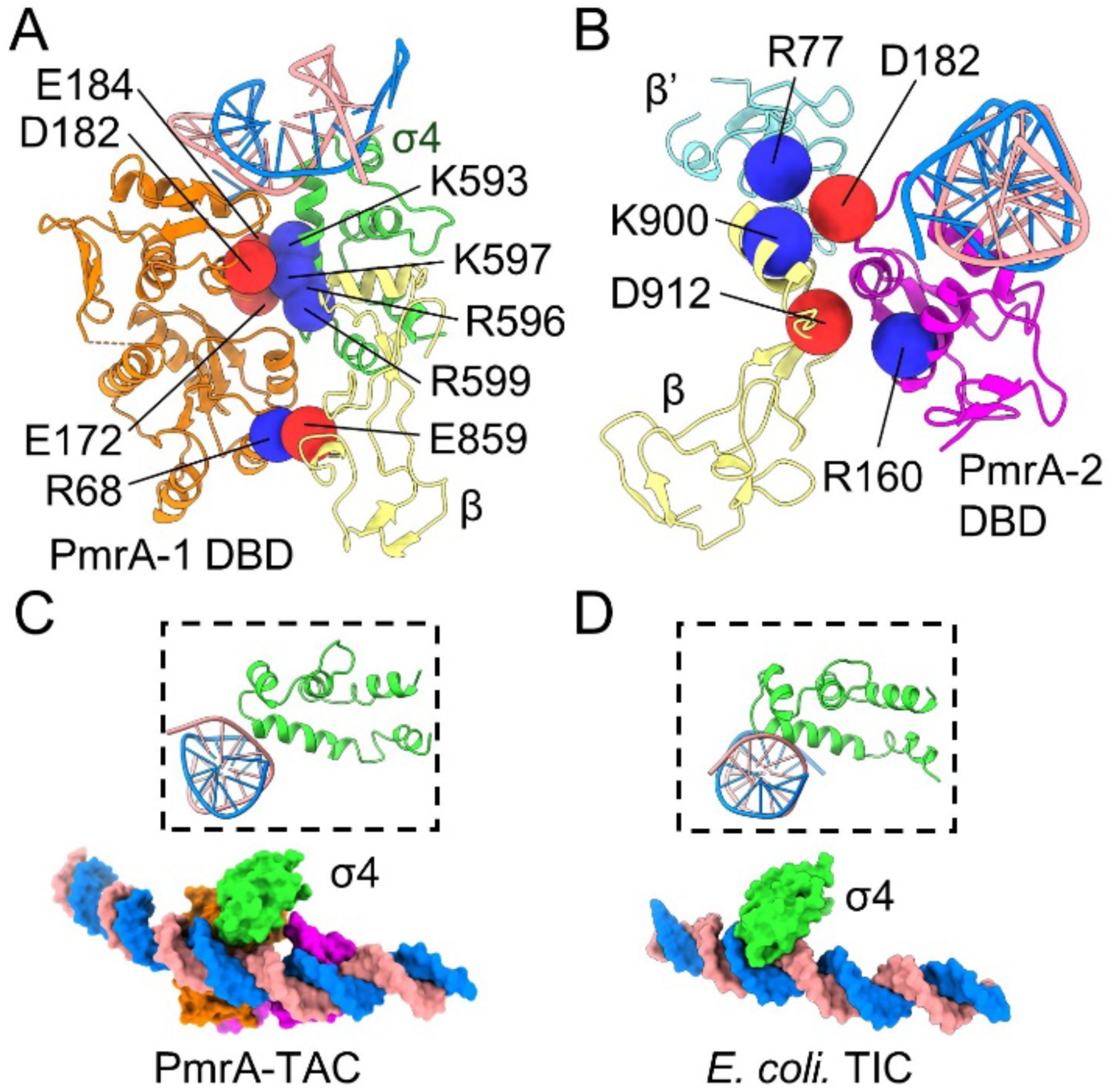
Analysis of the PmrA-RNAPH interface. Dimeric PmrA interacts with the double-stranded DNA, the σ4 domain of the σ70 cofactor, and the RNAP core β and β’ subunits. (A) The negatively-charged PmrA-1 DBD residues E172, D182, and E184 form electrostatic interactions with positively-charged residues K593, R596, K597, and R599 of the σ4 domain of the σ70 cofactor. Notably, residue R68 of the N-terminal receiver domain (REC) located close to E859 of the RNAP β subunit provides additional interactions to stabilize the complex. (B) PmrA-2 DBD that binds to the downstream DNA interacts electrostatically with the RNAP β and β’ subunits. (C) The interactions between the σ4 domain and DNA in the PmrA TAC and (D) the *E. coli* RNAP σ70 transcription initiation complex (TIC, PDB: 6CA0^21^). Due to binding of the two PmrA DBDs to the -35 position of the DNA, interaction between the DNA and the σ4 domain is weaker in the PmrA TAC than in the *E. coli* TIC. Lateral view (right top corner) showing that in the PmrA TAC, the σ4 domain is not inserted as deeply into the DNA major groove as it is in *E. coli* TIC without transcription factor.

Next, we examined the conformation of DNA in the PmrA TAC. We observed that the DNA near PmrA-binding regions shifted upon binding of that response regulator (Figure S3), so σ4 does not enter the DNA major groove as deeply as it does in the *E. coli* TIC in the absence of a transcription factor. In the PmrA TAC, the interface between σ4 and the DNA covers 127.7 Å^2^ (Figure 2C), i.e., much smaller than the interface of 504.4 Å^2^ between σ4 and the *-35 element* DNA in the *E. coli* TIC (Figure 2D). Thus, during activator-dependent transcription, both the recognition affinity and specificity of σ4 are weaker than those during activator-independent transcription because the weak interaction between σ4 and the promoter DNA is compensated for by its interaction with the activator, as shown in Figure 2.

### Upstream-shifted PmrA-DNA recognition

In our previous PmrA-DNA complex structure, the two DBDs bind to the DNA in a head-to-tail orientation, with the DNA recognition helix and the C-terminal β-hairpin (Cβ) inserted into the major and minor grooves, respectively, for DNA recognition, and the two N188 residues on the recognition helices specifically interacting with the “AA” bases of the imperfect repeated sequences “CTTAAT” and “CCTAAG” of the *pmra-box*. In the PmrA TAC, the two DBDs adopt a similar conformation to recognize DNA, but the two N188 residues lie in the upstream region far from those “AA” sequences (Figure 3A). We aligned the DNA structure in the previously generated PmrA-DNA complex with that of the PmrA TAC and found that the DBD domains in the two complexes indeed are not aligned with each other (Figure 3B). In the absence of the RNAPH, the DBDs bind to a major groove site “a”, where N188 contacts the “AA” bases. In the PmrA TAC, this site is occupied by σ4 and the β’ subunit of the RNAPH, so that both DBDs are shifted by RNAPH to the upstream site “b”. A lateral view (Figure 3C) clearly shows that the DBD binds to site “a” in the absence of the RNAPH and displays an 82º shift along the DNA major groove to site “b” upon interaction with the RNAPH, representing an upstream shift of approximately three base pairs. In this shifted DNA-binding position, the interactions between PmrA and DNA are likely weaker than those identified for PmrA-DNA alone. This result correlates well with our initial cryo-EM map of the PmrA TAC, which shows a dynamic TAC conformation with a clear density map for the RNAPH, but weak signals for the upstream DNA and PmrA.

**Figure 3.**
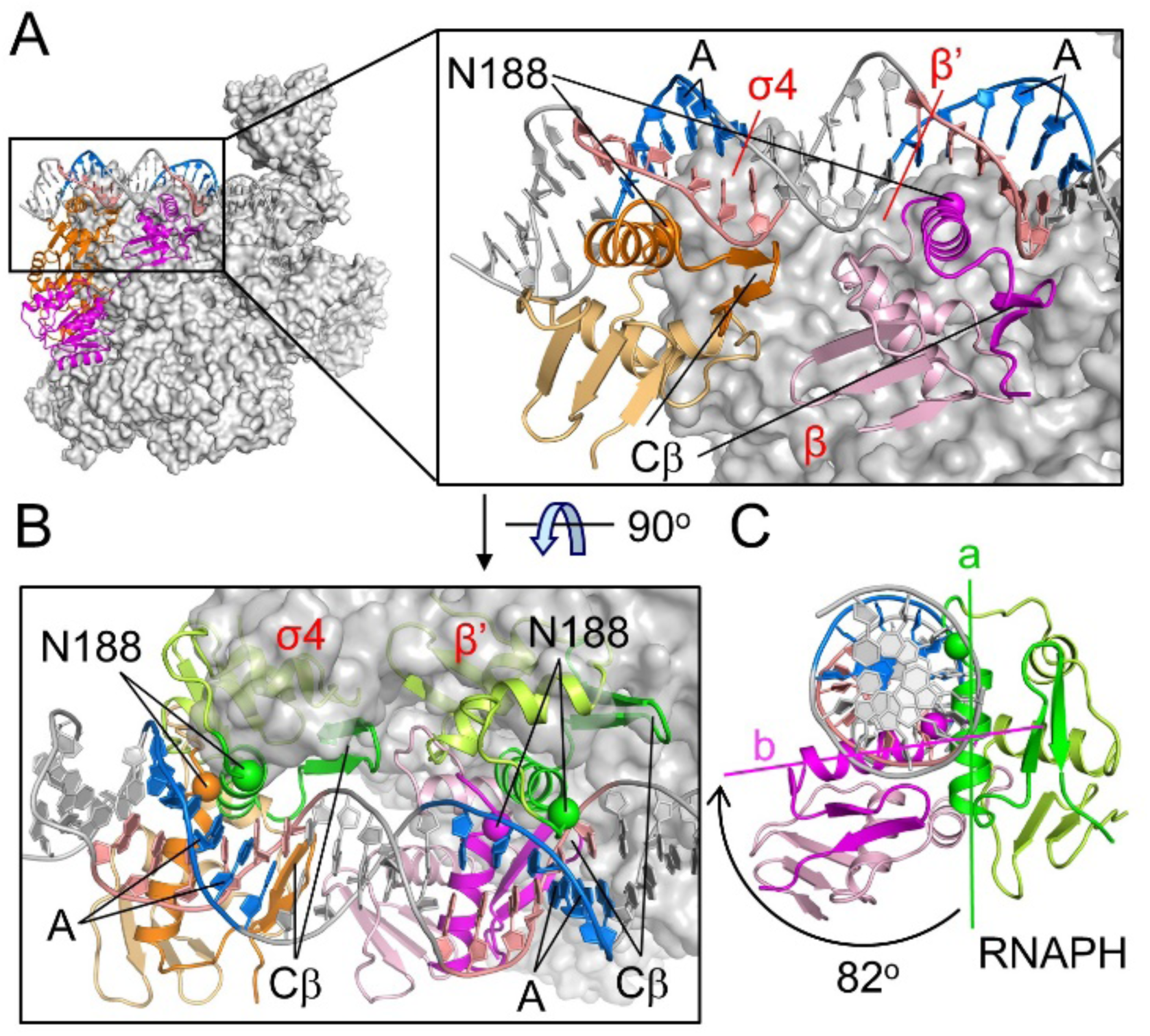
Upstream-shifted PmrA-DNA interaction in the PmrA TAC. (A) In the PmrA TAC, the two DBDs of PmrA contact DNA, with the DNA recognition helix and the C-terminal β hairpin (labeled as Cβ) inserted into the major and minor grooves, respectively. Residue N188 on the recognition helix (shown as a sphere) specifically contacts the “AA” bases of the imperfect repeat sequences “CTTAAT” and “CCTAAG” in the PmrA-DNA complex structure. However, in the PmrA TAC, the two N188 residues are far from those “AA” bases. (B) The 90° rotated view shows the DBD domains in the PmrA-DNA complex structure (green for the recognition helix and Cβ, and others in light green) and in the PmrA TAC (magenta and orange for the recognition helix and Cβ, and others in light colors) with structural alignment on the DNA. This view clearly shows that DBD binding regions in the PmrA-DNA complex without the RNAPH are occupied by the σ4 domain and β’ subunit of the RNAPH in the PmrA TAC. (C) Lateral view showing that the recognition helix in the DBD binds to position “a” in the absence of the RNAPH (colored green), and is shifted upstream by 82º along the DNA major groove to position “b” in the PmrA TAC (colored magenta).

### PmrA residues responsible for promoter and RNAPH interaction are also important in transcription

In our PmrA TAC structure, the RNAPH contacts PmrA via several electrostatic interactions, but also shifts PmrA to the upstream DNA sequence, resulting in weaker PmrA-DNA interactions. To understand how those interactions contribute to transcriptional activation by PmrA, we performed *in vivo* β-galactosidase reporter gene expression assays (Figure 4A). Mutating PmrA residue N188 to alanine (N188A) to abolish DNA recognition activity^14^ also eliminated its transcriptional activity, implying that DNA recognition is essential for PmrA-mediated transcriptional activation. Interestingly, alanine replacement of any one of the four residues involved in RNAPH interaction (R160, E172, D182 and E184) enhanced gene expression. Moreover, replacement of any of the three negatively-charged residues with a positively-charged lysine (E172K, D182K and E184K) elicited even higher gene expression. Control experiments using alanine replacement of either K164 or H170, which are not involved in RNAPH interaction, did not alter expression or transcriptional activity (Figure 4A).

**Figure 4.**
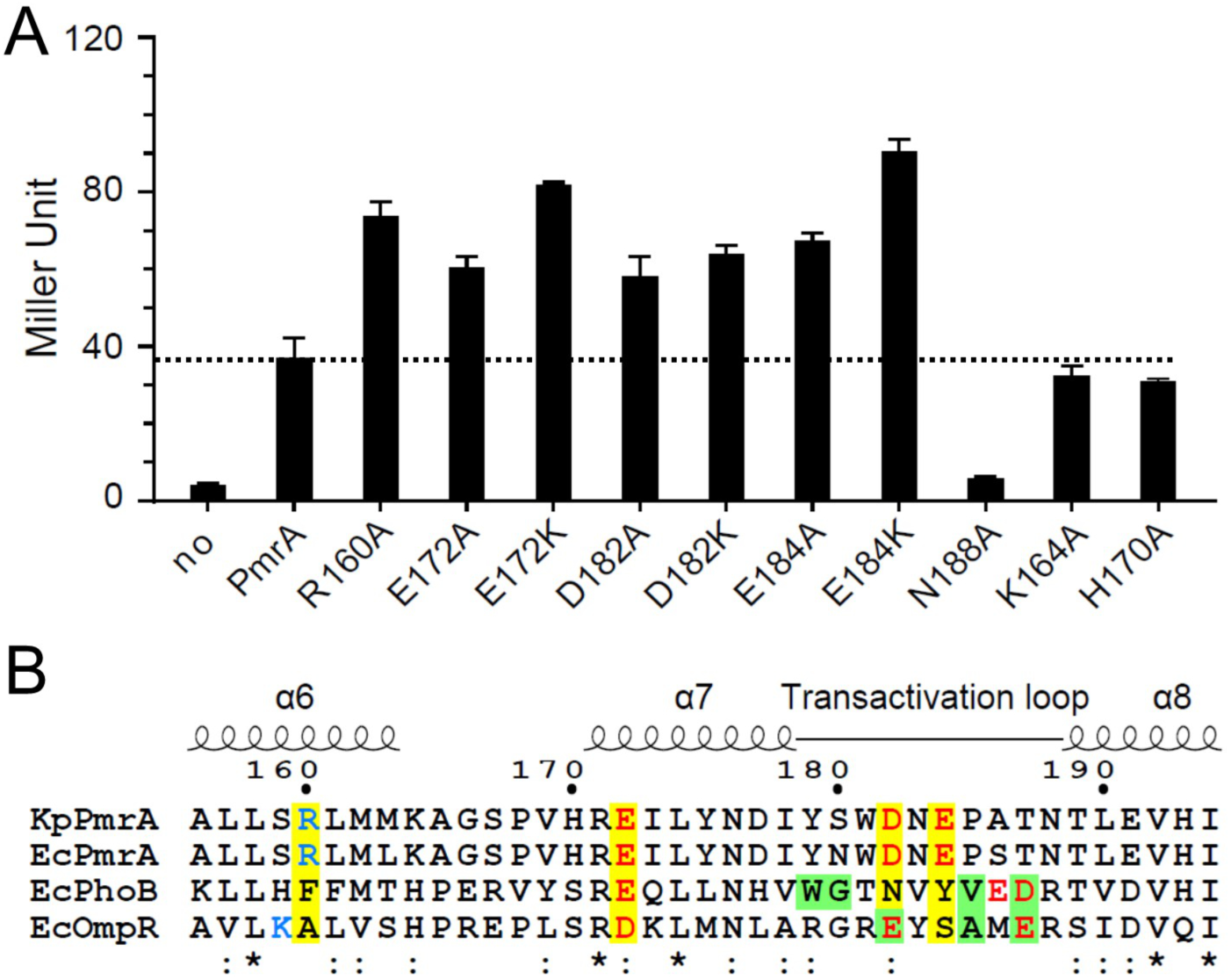
Residues involved in transcriptional activation. (A) *In vivo* β-galactosidase reporter assay in *K. pneumoniae* carrying plasmids that expresses PmrA or its variants plus the reporter plasmid hosting the *pbgP* promoter in front of the *lacZ* gene. Expression of PmrA or its variants was induced by isopropyl-β-D-thiogalactoside (IPTG, 1 mM/ml). Assay for PmrA without IPTG addition is labeled as “no”. Results are expressed as Miller Units. All experiments were performed in triplicate. Error bars represent standard deviation. Horizontal dashed line indicate the values obtained from PmrA. (B) Sequence alignment of *K. pneumoniae* PmrA, and PmrA, PhoB and OmpR from *E. coli. K. pneumoniae* PmrA residues involved in RNAPH interactions are highlighted with a yellow background and are colored in blue and red for positive and negative charges, respectively. Residues for which replacement abolishes transcriptional activation by *E. coli* PhoB and OmpR are shown with a green background. Fully conserved or similar residues are denoted * or :, respectively.

### Comparison of PmrA with PhoB and other transcription factors

Our structure-function analysis indicates that residues R160, E172, D182 and E184 that are involved in RNAPH interactions are important to PmrA-dependent transcriptional activation. Amino acid sequence alignment of *K. pneumoniae* PmrA with *E. coli* PmrA, PhoB and OmpR reveals conserved or similar residues at proximal positions in all those response regulators (Figure 4B). Previous mutational analyses have shown that replacement of residues on the transactivation loop abolishes transcriptional activation by *E. coli* PhoB and OmpR (residues with a green background in Figure 4B) ^15,19,24^. However, in *K. pneumonia* PmrA, variants in which RNAPH-interacting residues had been altered (in the current study, two residues on the transactivation loop) resulted in elevated transcriptional activity.

To understand why substitution of these two residues on the transactivation loop exert a differential effect on transcriptional activation, we compared the promoter sequences of PmrA, PhoB and two other well-studied transcription factors, i.e., TAP (a homolog of *E. coli* cyclic receptor protein CAP), and EcmrR (a MerR family regulator) (Figure 5A). We excluded OmpR promoters from our analysis because OmpR binds to DNA upstream of the *-35 element* and interacts with the C-terminal domain of the α subunit^25^. As shown in Figure 5A, it is clear that EcmrR contacts the downstream region between the *-35* and *-10 elements*, TAP recognizes the upstream region of the *-35 element*, and both PmrA and PhoB bind to an intervening region. EcmrR and TAP contact different regions on the promoters and activate transcription via different mechanisms. In the class-II transcriptional activation elicited by *E. coli* CAP^26^ or by *Thermus thermophilus* TAP^27^, the transcription factor interacts with the RNAP β’ subunit and the σ4 domain of the σ70 cofactor to stabilize the open complex (Figure 5B). RNAPH recruitment is dominant in this mechanism, so substitution of any residues involved in RNAPH interactions inhibits transcriptional activation. In terms of transcriptional activation by the MerR family regulator, the EcmrR TAC structure^28^ reveals that the RNAPH and EcmrR recognize the promoter DNA from opposite sides, without any interaction, but EcmrR binding induces significant kinks in DNA that shortens the non-optimal *−35/−10 element* spacers for optimal promoter recognition by the RNAPH (Figure 5C). Thus, the MerR family of regulators activate transcription by distorting DNA in an RNAPH-contact-independent manner.

**Figure 5.**
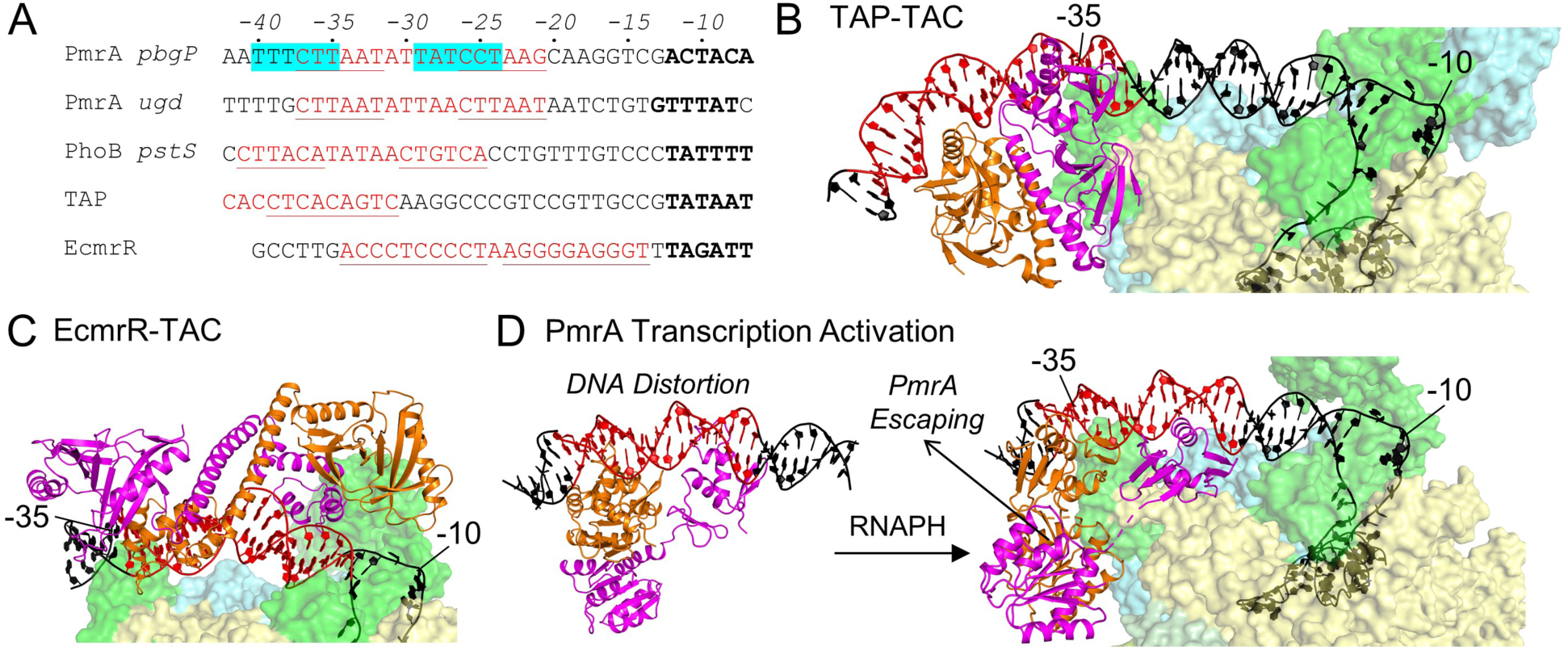
The complex mechanism of PmrA-modulated transcriptional activation. (A) Alignment of the *pbgP* and *ugd* (for PmrA), and *pstS* (for PhoB) promoter sequences, as well as TAP (a *Thermus thermophilus* homolog of *E. coli* CAP) and EcmrR (MerR family regulator) transcription factors. The transcription factor recognition sites are marked in red and underlined. Shifted PmrA-binding sites in the PmrA TAC are highlighted with a cyan background. The *-10 elements* are marked in bold. The TAP-TAC (PDB: 5i2d) and EcmrR-TAC (PDB: 6xl5) structures are shown in (B) and (C), respectively. Both transcription factors adopt a dimeric conformation and are colored in orange and magenta. (D) The PmrA-promoter DNA complex and PmrA TAC structures are shown in the left and right panels, respectively. From B to D, the DNA is colored as in (A) and color-coding of the transcription factors and RNAPH are as in Fig. 1(A). The -35 and -10 positions are also indicated.

In the PhoB-recognizing *pstS* promoter^19^, the PhoB binding sequence spans from the -25 to -41 positions, which is four base pairs upstream of the *pmra-box* and similar to the PmrA-binding position identified for the PmrA TAC. Hence, the RNAPH can directly contact PhoB on the promoter, in a scenario similar to the mechanism of class-II transcriptional activation with dominant RNAPH recruitment, potentially explaining why the mutants in which residues of the PhoB transactivation loop were replaced lacked transcriptional activation capability.

In the PmrA-regulated *pbgP* and *ugd* promoters, the PmrA dimer specifically recognizes two “TAA” sequences in the *pmra-box*, which bends the promoter DNA (Figure 5D, left)^14,22^. PmrA-binding and consequent DNA distortion are necessary for RNAPH recognition, as evidenced by the N188A mutation abolishing transcriptional activity. However, the PmrA binding sites are also elements of the RNAPH recognition region, so the PmrA dimer shifts upstream upon RNAPH binding. The shifted PmrA dimer can then contact RNAPH via several electrostatic interactions, but also recognizes the DNA through weaker interactions, thus enabling PmrA to escape from the promoter DNA, representing the next step in the transition from TAC to the elongation complex (Figure 5D, right). Substitutions of RNAPH-interacting residues in PmrA result in elevated, and not diminished transcriptional activity, suggesting that PmrA escaping from the promoter is more important than RNAPH recruitment. Notably, the PmrA E172K, D182K and E184K mutations all elicited the greatest effect on gene expression because these mutated residues repulse the positively-charged residues on RNAPH, so those variants can be more easily pushed away from the DNA. Taken together, DNA distortion caused by PmrA binding to the *pbgP* and *ugd* promoters and subsequent PmrA escape from those promoters are important for their transcriptional activation.

## CONCLUSION

In summary, we reveal dynamic coupling among subunits of the TAC structure containing a dimer of full-length OmpR/PhoB-family response regulator PmrA, a RNAPH, and the *pbgP* promoter. Our TAC structure shows that PmrA contacts RNAPH via several electrostatic interactions, most notably between the two DBD domains and the σ4, β, β’ subunits of the RNAPH, implying that RNAPH recruitment is essential for transcription. However, compared to the PmrA-DNA complex structure, binding of the RNAPH shifts both DBD domains of the TAC upstream by approximately three base pairs, resulting in weaker interaction with the promoter DNA. Moreover, we found that PmrA variants hosting mutations of RNAPH-interacting residues, which should display weaker interaction with the RNAPH, exhibited enhanced transcriptional activity. Taken together, we propose that PmrA exit<u>s</u> from the complex to allow progression into the elongation state is important for transcriptional activation. In addition, the PmrA-DNA complex structure revealed a bent DNA conformation, with alanine substitution of PmrA DNA-recognition residue N188 nearly abolishing transcriptional activation, indicating that the DNA distortion induced upon binding of PmrA also is important for this activity. Comparing promoter protein binding sites, activation by PhoB is similar to the mechanism of simple RNAP recruitment identified for class-II transcriptional activation by the *E. coli* CAP. In contrast, here we have shown that the mechanism of PmrA-modulated transcriptional activation is more complex than that of simple RNAP recruitment.

## Acknowledgments

We thank the Ministry of Science and Technology of Taiwan for support to K.-P.W. (MOST 110-2311-B-001-046) and C.C. (MOST 108-2311-B-001-016-MY3). K.-P.W. received a career development award (AS-CDA-110-L03) from Academia Sinica (AS). We appreciate Dr. Meng-Ru Ho (Biophysics Instrumentation Laboratory, Institute of Biological Chemistry, AS) for assistance with and valuable discussions on the biophysical characterization of PmrA-related complexes. The cryo-EM experiments were performed at the Academia Sinica Cryo-EM Facility (ASCEM). ASCEM is jointly supported by the Academia Core Facility and Innovative Instrument Project (AS-CFII-108-110) and Taiwan Protein Project (AS-KPQ-109-TPP2).

