## Supplemental information for "Structural basis of transcriptional activation by the OmpR/PhoB-family response regulator PmrA"

SI Materials and Methods
SI Figures S1-S3
SI References

### Materials and methods

#### Preparation of the PmrA-dependent TAC

The DNA fragment encoding *K. pneumoniae* PmrA hosting the W181G/I220D double-substitution, together with an extra Met residue at the N-terminus and an additional LEHHHHHH tag at the C-terminus, was cloned into a pET-29b(+) (Novagen) vector and transferred into *E. coli* strain BL21(DE3). Recombinant PmrA was expressed and purified as described[1]. A synthetic promoter comprising the upstream AT-rich sequence and the *pmra-box* from the *K. pneumoniae pbgP* promoter, a consensus *-10 element*, a discriminator element, a transcription bubble, and a 14-bp downstream extension was prepared by mixing an equal amount of two strands, heating to 95 °C for 30 min and cooling slowly to room temperature. The polycistronic vector for expressing *E. coli* RNAP was obtained from Addgene (plasmid #104398)[2]. The DNA fragment encoding the *E. coli* $\sigma^{70}$ cofactor was directly amplified from *E. coli* DNA and cloned into a pGEX-4T1 vector with a modified Tobacco etch virus (TEV) protease cleavage sequence. The two vectors were transferred into *E. coli* strain BL21(DE3) for separate expression. Cells were grown in LB broth at 37 °C. When OD $A_{600}$ reached 0.6, protein expression was induced by means of 1 mM IPTG at 28 °C overnight. Cells were harvested, resuspended in lysis buffer (50 mM Tris, 500 mM NaCl, 5% glycerol at pH 7.3), and homogenized by using a microfluidizer. After lysis, cell debris was removed by centrifuging at 12000 rpm for 30 min. The cell lysates containing RNAP and $\sigma^{70}$ cofactor were mixed for 1 h and purified by using Ni-affinity chromatography and a GSTrap FF column. The RNAP $\sigma^{70}$-holoenzyme (RNAPH) was eluted from the GSTrap FF column by using buffer containing 50 mM Tris, 200 mM NaCl, 20 mM reduced-glutathione at pH 8.0. The GST-tag was removed by using the TEV protease. The sample of RNAPH was further purified by size-exclusion chromatography and examined on a Coomassie blue-stained SDS polyacrylamide gel (Fig. S1). The PmrA-dependent TAC was assembled by incubating the RNAPH with a 2-fold molar excess of the preformed promoter DNA and PmrA in final buffer (20 mM HEPES, 100 mM NaCl, 10 mM $MgCl_2$, 200 μM $ZnCl_2$, pH 7.0).

#### Cryo-EM sample, data collection, data process and structure building

The purified PmrA TAC was thawed on ice from -80 ˚C and diluted to 0.3-1 mg/ml for cryo-EM grid preparation. The protein complex was applied to Quantifoil holey carbon grids (Cu, 200 mesh, R2/1 μm) by using a Vitrobot Mark IV system (Thermo) with 4-s blotting time at 100% humidity and 4 ˚C. Grids were immediately frozen by using liquid ethane and stored in a liquid nitrogen reservoir. Cryo-EM data for the PmrA TAC were collected on a 300 keV Titan Krios microscope equipped with a Gatan K2 detector. The slit width was 20 ev. EPU 1.2 was used for automatic data collection with three exposures per hole. In total, 6132 60-frame micrographs in counting mode were acquired at a physical pixel size of 0.822 Å and a nominal magnification of 160,000. The total electron dose and desired defocus ranges were 57.1 e-/Å$^2$ and -1.0, -1.5, -2.0, and -2.2 μm, respectively.

Cryo-EM data were processed by using cryoSPARC 2.12 and, later, 3.2[3]. Movie alignment was performed by full-frame motion or patch-motion correction. The contrast transfer function was estimated by using Patch CTF in cryoSPARC. Particles with 150- to 180-Å blob diameters were extracted from 150 micrographs for 2D classification. The well-aligned 2D averages were used for template particle selection in cryoSPARC. Through iterations of 2D classifications, 424,370 particles were finally selected for 3D map refinement. Applying 3D heterogeneity to separate the PmrA TAC complex from the RNAP and RNAPH was unsuccessful because the EM density of the PmrA region proved consistently fragile. Therefore, the 424,370 particles were used for 3D homogeneous refinement in cryoSPARC, followed by 3D variability analysis (3DVA) with a filter resolution of 5 Å. We selected a subgroup of five 3DVA clusters containing 55,568 particles of the PmrA TAC complex for non-uniform refinement to 3.03 Å (Fig. 1B). The map was used to build the RNAPH structure. By analyzing the 3DVA series mode (20 frames per series), we revealed a substantial density for PmrA in the cryo-EM map, which was used for further model building focused on PmrA and PmrA-associated interactions.

Cryo-EM structures were built using existing template structures for *E. coli* RNAPH, including PDB IDs 6CA0, 6BC6, and 6B6H. The 3.03-Å map was first used to remodel RNAP core subunits and $\sigma^{70}$ cofactor by means of the real-space refinement function in Phenix 1.8[4]. Subsequently, the DNA sequences of this study were manually replaced into the structure by using Coot 0.8.2[5]. The DBD and REC domains of PmrA were separately docked into the 3DVA enhanced map, followed by real-space refinement. The final model was manually inspected and verified in Coot.

**β-Galactosidase reporter assay.**

The upstream region (~500 bp) of the *K. pneumoniae pbgP* gene was PCR-amplified and inserted in front of the *lacZ* gene in the placZ15 plasmid to obtain the reporter plasmid, which was then transformed by conjugation into *K. pneumoniae* CG43S3-*ΔlacZ* strain. Plasmids expressing PmrA or its variants were then transformed into the *K. pneumoniae* cells carrying that reporter plasmid. The cells were cultured at 37 °C in LB medium containing kanamycin (50 mg/ml) and chloramphenicol (34 mg/ml) for 30 mins, before adding isopropyl-β-D-thiogalactoside (1 mM/ml) for PmrA or variant expression for 5 h and then harvesting by centrifugation. β-Galactosidase activity in the supernatant was measured at $OD_{420}$, and values are expressed in Miller units ($1{,}000 \times OD_{420}/(T \times V \times A_{600})$), where T and V are reaction time (minutes) and volume of culture (ml), respectively. The average values and standard errors from triplicate measurements are plotted.

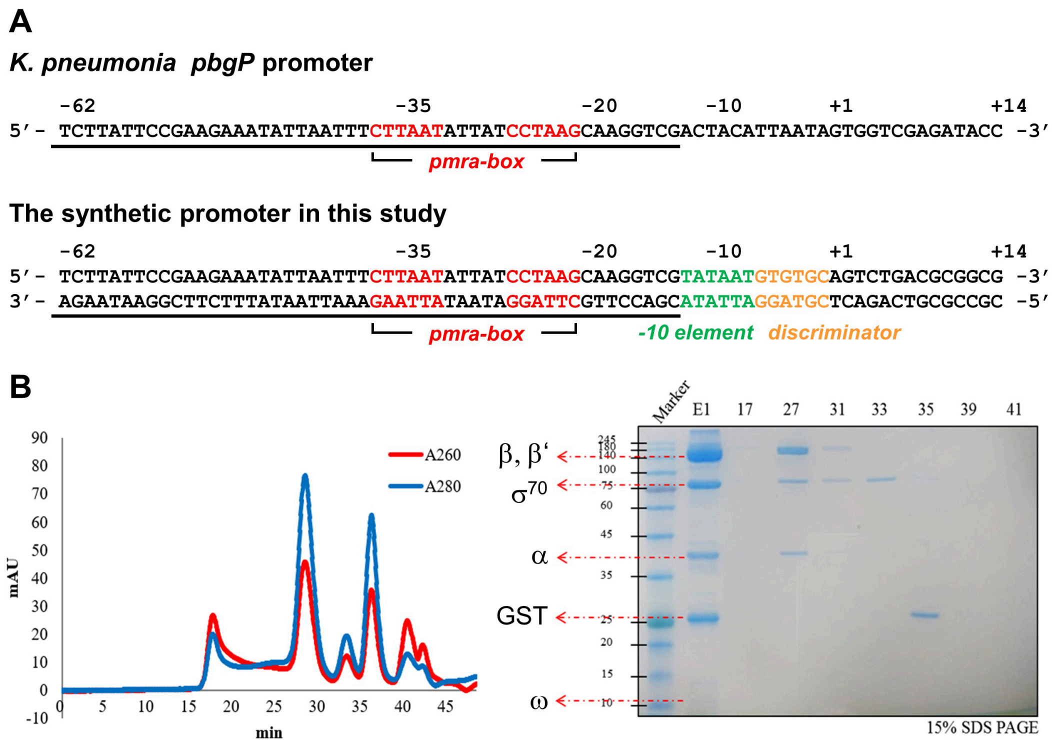


**Fig. S1: DNA promoters and preparation of the RNA polymerase $\sigma^{70}$-holoenzyme (RNAPH).**

(A) DNA sequences of the *K. pneumonia pbgP* promoter and the synthetic promoter for the PmrA TAC. The *pmra-box*, *-10 element* and *discriminator element* are in red, green, and orange, respectively. (B) Size exclusion chromatography (SEC) showing the absorption profiles at wavelengths 260 and 280 for the RNAPH separated by a Superose6 10/300 GL column (GE Healthcare). The SEC fractions were analyzed by 15% SDS-PAGE. Lane E1 represents the sample before SEC. Fraction 27 shows co-elution of RNAP subunits and the $\sigma^{70}$ factor.

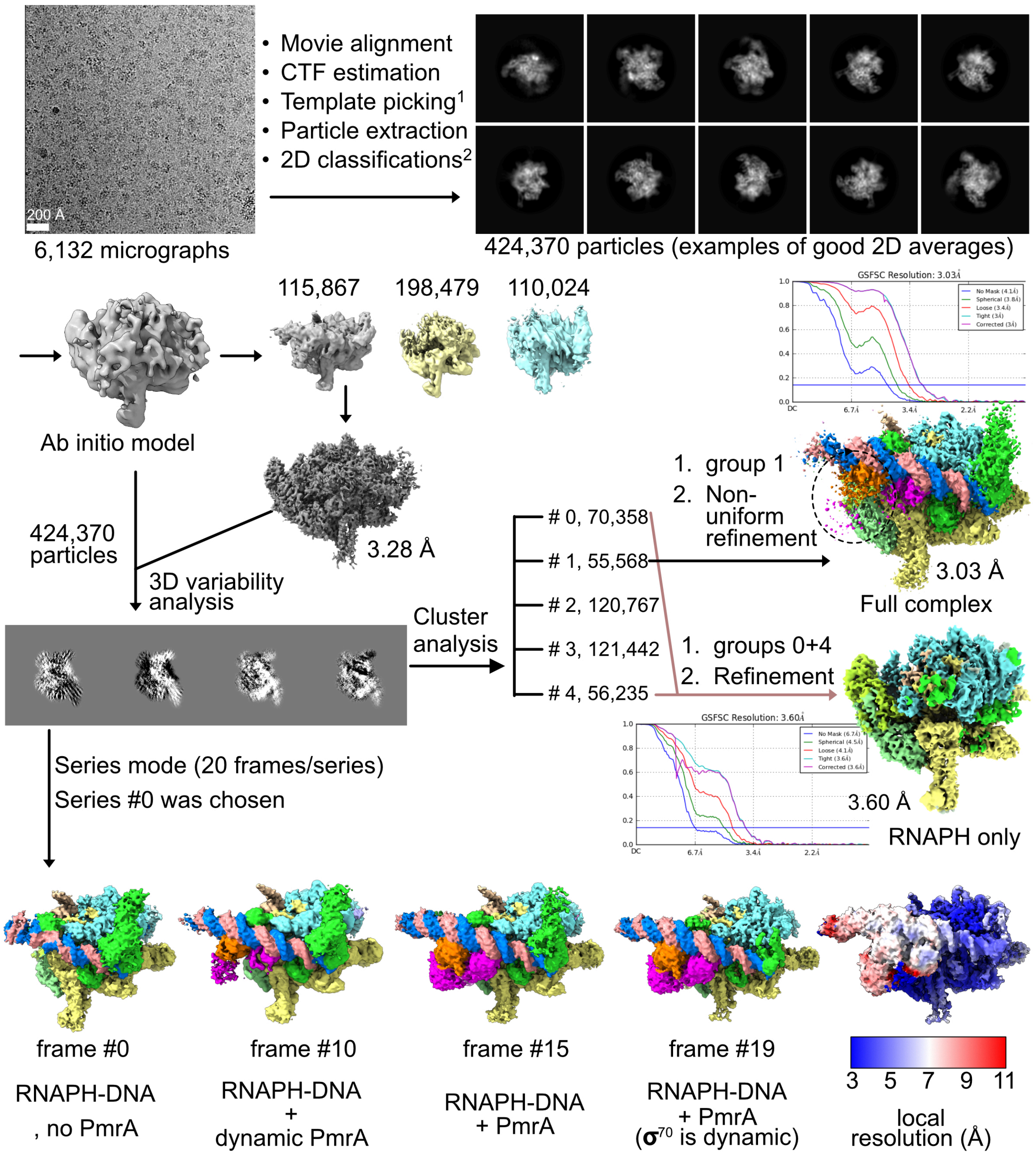


1: using selected 2D classes from 50-micrograph data, 1,113,892 particles picked

2: 100 groups

**Fig. S2: Workflow for cryo-EM single particle reconstruction of the RNAPH-DNA-PmrA complex.**

The entire dataset was processed initially using cryoSPARC 2.12 and then version 3.2 for 3D variability analysis (3DVA). The raw 60-frame movies were aligned and contrast transfer function (CTF) was estimated by using patch motion and patch CTF, respectively. A total of 50 micrographs was used to select blob particles, followed by 2D classifications. Appropriate 2D averages were then used as templates to identify RNAP-related particles automatically from 6132 images. Approximately, 424,000 particles were selected for 3D map calculations. Heterogeneous refinement resulted in a complex map, but the densities for DNA and PmrA were extremely weak, indicative of dynamic transitions. Therefore, we adopted a 3DVA approach to identify four distinct modes of the TAC complex. Analyses of clusters and series based on the four 3DVA modes were performed separately to reveal the whole-complex map (55,568 particles at 3.03 Å) and dynamic PmrA binding. We also reconstructed an RNAPH map at 3.6 Å, but the preferred orientation limited further model building. In the series mode analysis (20 frames), both RNAPH-DNA and RNAPH-DNA-PmrA states were characterized from frame numbers 10 to 19. Frame #19 was used to build the DNA-PmrA complex. The local resolution estimation colored on frame #19 is also displayed showing a dynamic RNAPH-bound PmrA.

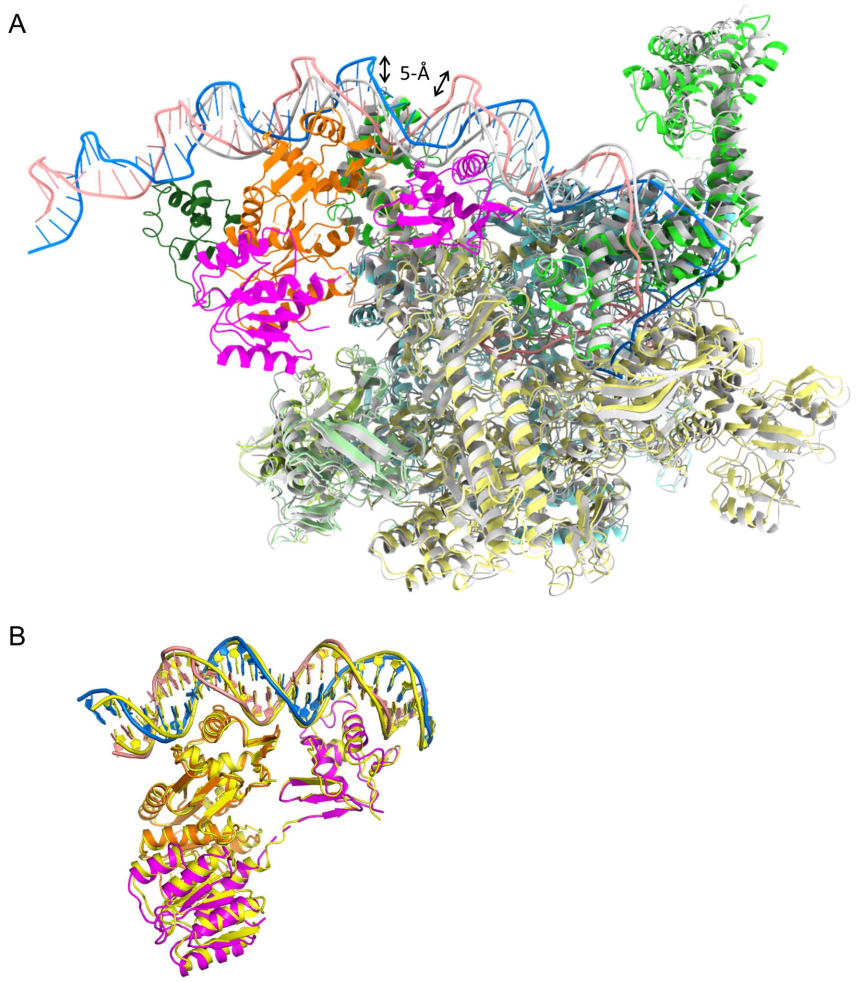


**Fig. S3: Structural comparisons with the *E. coli* RNAP σ70 transcription initiation complex (TIC) and the PmrA-DNA complex.**

(A) Superimposition of the PmrA TAC complex with the *E. coli* TIC (PDB ID: 4YLN) centered on the RNAP. Color codes of the PmrA TAC are consistent with those of Fig. 1, with the *E. coli* TIC in gray. (B) Superimposition of PmrA from the PmrA TAC (orange for PmrA-1 and pink for PmrA-2) with PmrA in complex with DNA alone (in yellow, PDB ID: 4S04).
